# Genomic Erosion Through the Lens of Comparative Genomics

**DOI:** 10.1101/2025.03.28.645899

**Authors:** Xuejing Wang, Claudia Fontsere, Ximena Alva Caballero, Sascha Dreyer Nielsen, Jim Groombridge, Bengt Hansson, Cock van Oosterhout, Carolina Pacheco, Hernán E. Morales

**Author notes:** These authors contributed equally (Xuejing Wang, Claudia Fontsere, Carolina Pacheco, Hernán E. Morales).

## Abstract

Loss of genetic diversity threatens species survival, yet the dynamics of such loss and species’ responses thereof can vary widely depending on their evolutionary histories, life-history traits and demographic trajectories. Comparative genomics offers a powerful framework to explore the dynamics of genomic erosion across species. Here, we analysed the genomes of three species — the Mauritius parakeet, the Mauritius kestrel, and the pink pigeon — that experienced extreme and well-documented population bottlenecks. We compared them to 36 species spanning the avian phylogeny, with varied IUCN Red List statuses to investigate the genomic consequences of their demographic collapses. For each species, we assessed nucleotide diversity, genetic load, and runs of homozygosity (ROH), alongside genome synteny and transposable elements. We found a negative correlation between nucleotide diversity and ROH, but neither metric was a good predictor of the species’ Red List status. Rather, the population effective to census size ratio showed a strong correlation to Red List status. Moreover, species with larger historical effective population sizes showed greater heterozygosity but carried a higher heterozygous load, highlighting the importance of historical demography to assess species vulnerability to genomic erosion. We found significant differences in homozygous load between taxonomic groups of our target species, possibly due to differences in life-history traits and demographic histories. Genome structure analyses revealed differences in transposable elements and genomic rearrangements between groups, suggesting their potential role in shaping genome architecture and adaptive potential across species. Our findings underscore the value of multispecies comparisons in understanding the evolutionary dynamics of genomic erosion and its relevance for biodiversity conservation.

## Introduction

Genetic diversity is an essential component of a species’ ability to adapt and persist under changing environmental conditions (Spielman et al. 2004; Allendorf et al. 2013; Kardos et al. 2021). For small or isolated populations, maintaining genetic diversity is particularly challenging, as reduced effective population size (Ne) diminishes the efficacy of natural selection, intensifies genetic drift, and leads to increased inbreeding. These processes ultimately lead to genomic erosion, often characterised by the loss of genetic diversity and an increased accumulation or fixation of deleterious mutations. As inbreeding becomes more frequent, the exposure of recessive deleterious mutations exacerbates these effects, culminating in inbreeding depression (Charlesworth and Willis 2009; Blomqvist et al. 2010; Hasselgren et al. 2021). These processes collectively diminish fitness and adaptive potential, exacerbating vulnerability to environmental changes and threatening the long-term persistence of the population (Blomqvist et al. 2010; Hasselgren et al. 2021; Jackson et al. 2022; van Oosterhout et al. 2022; Jeon et al. 2024). Additionally, even populations that have partially recovered demographically bear the genetic legacy of past bottlenecks, known as “drift debt”, which manifests as a time lag between demographic change and loss of genome-wide variation (Dussex et al. 2023; Gilroy et al. 2017; Pinto et al. 2024). Recent analyses have shown that species are losing genetic diversity worldwide (Exposito-Alonso et al. 2022; Shaw et al. 2025), highlighting the critical need for understanding genomic erosion in biodiversity conservation.

With the continuous and rapid production of genomic data for wild species worldwide, conservation genomics can now take advantage of high-resolution tools to assess genetic diversity, genetic load and structural variation (Lewin et al. 2018; Wright et al. 2020; van Oosterhout et al. 2022; Theissinger et al. 2023). For species at risk of extinction, such insights are critical for guiding conservation interventions aimed at reducing genetic load and enhancing population viability (e.g., vaquita (Morin et al. 2021), kākāpō (Dussex et al. 2021), pink pigeon (Speak et al. 2024)). Additionally, the use of genomic resources across multiple species with diverse evolutionary histories, demographic trajectories, and conservation status within a comparative genomic framework offers valuable insights (Grueber 2015). This approach can elucidate how these factors interact with genomic traits such as genetic diversity, genetic load, or structural variations to influence species’ long-term viability and extinction risk.

The echo parakeet (*Alexandrinus [Psittacula] eques*), Mauritius kestrel (*Falco punctatus*), and pink pigeon (*Nesoenas mayeri*) exemplify how species can recover demographically from near extinction but remain genetically imperilled. These birds, endemic to Mauritius—a biodiversity hotspot that has witnessed over 100 species extinctions in recent centuries (Florens 2013)— experienced some of the most extreme population bottlenecks ever recorded in wild populations but recovered thanks to targeted conservation efforts (Jones and Swinnerton 1997). Only four Mauritius kestrels remained by 1974, the pink pigeon declined to 10 individuals by 1990, and the Mauritius parakeet to just 20 individuals by 1986 (Jones and Swinnerton 1997; Jones 2013; Jones et al. 2013). Conservation programs, including captive breeding, supplementary feeding, habitat restoration, and predator control, facilitated their demographic recovery to current free-living adult population sizes of approximately 500 pink pigeons, 250 Mauritius kestrels and 650 echo parakeets (Jones and Swinnerton 1997; Jones 2010; Nicoll et al. 2021) (Figure 1D). Despite successful demographic recoveries, the legacy of the extreme historical population collapses can jeopardise their long-term viability, as genetic diversity in these species continues to decline due to the accrued drift debt (Tollington et al. 2013; Jackson et al. 2022). These species are also at risk of accumulating an increased genetic load of deleterious mutations, as has been shown in the pink pigeon (Jackson et al. 2022). Beyond genetic challenges, ecological pressures such as habitat loss and degradation persist, compounded by threats like emerging infectious diseases, such as the Psittacine Beak and Feather Disease (PBFD) in the Mauritius parakeet, can affect individuals fitness and population viability(Tollington et al. 2015).

**Figure 1.**
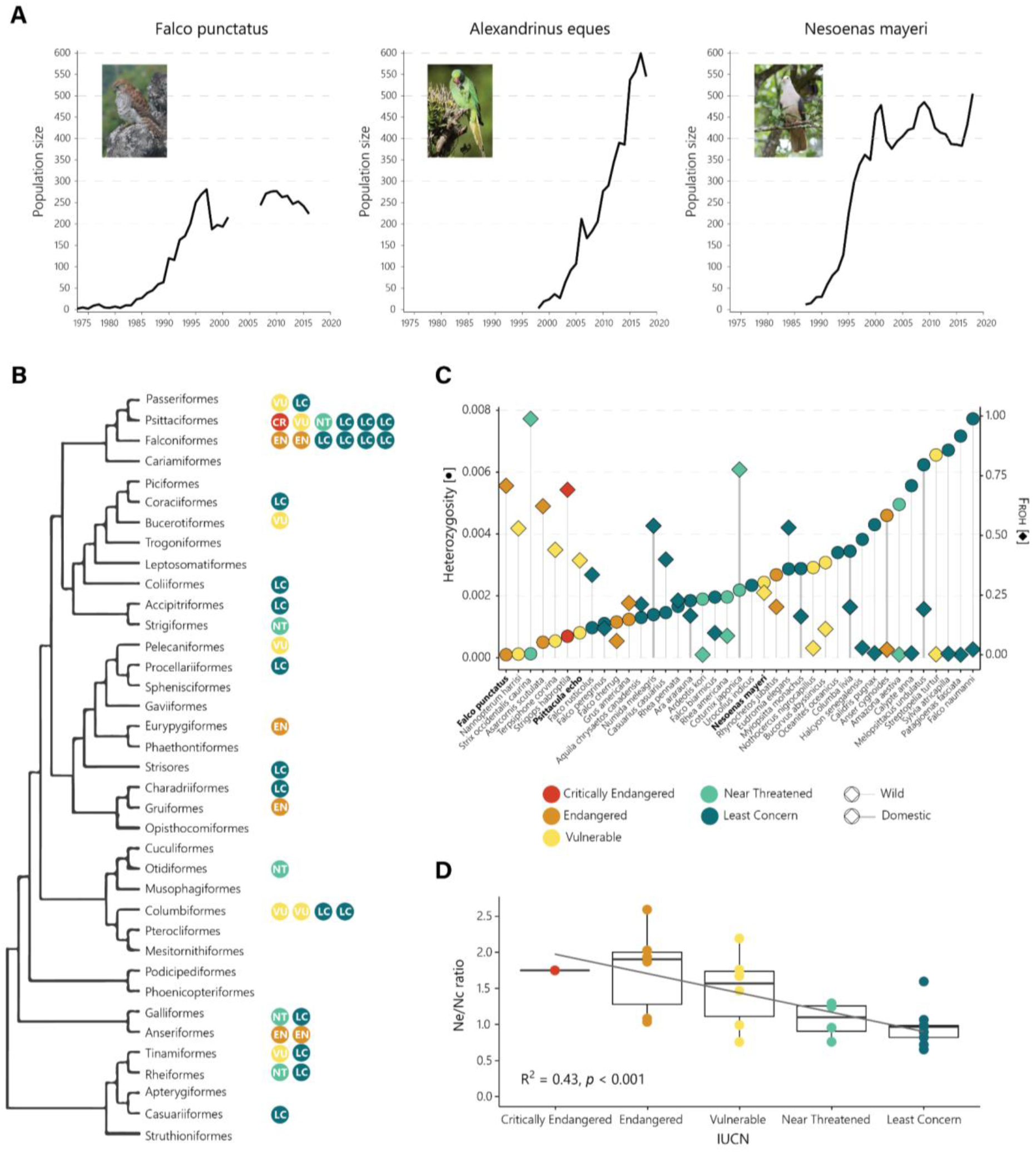
Demographic trajectory of the three Mauritius species and phylogenetic, genetic diversity, and Ne/Nc ratio distribution of the 39 species used in this study. **A** Demographic trajectory of the wild population derived from field monitoring of adult individuals over time. Note that the three species have been monitored in different ways so presented trends are approximations of their total numbers. **B** Phylogenetic tree topology, adapted from Stiller et al. (2024). Each circle represents a sampled species within its respective order. The colour and initials indicate the IUCN Red List category of each species. **C** Genome-wide heterozygosity (circles) and runs of homozygosity-based inbreeding coefficient (F_ROH_; diamonds) for each species. Colour coding corresponds to IUCN Red List categories. Domestic species are highlighted with a thicker vertical line. **D** Correlation between the log-transformed Ne/Nc ratio (Log(Ne)/Log(Nc)) and IUCN Red List categories. The effective population size (Ne) was estimated as the harmonic mean of PSMC values from 10 kya to 100 kya, whereas the census population size (Nc) was obtained from IUCN Red List data. Photo credits: Samantha Cartwright for the Mauritius kestrel (*Falco punctatus*), Jacques de Speville for the Mauritius parakeet (*Alexandrinus eques*), and Gregory Guida for the pink pigeon (*Nesoenas mayeri*).

This study aims to leverage the power of comparative genomics to explore the interplay between genome-wide diversity, genetic load, demographic history, and conservation status across a diverse set of avian species. Using recently generated high-quality, chromosome-level reference genomes for these three Mauritian birds, we compared them to a set of bird species spanning the avian phylogeny with distinct evolutionary histories and conservation statuses. By narrowing our focus to species within the same order as the Mauritius ones, we further aim to investigate potential group-specific differences in genomic metrics. This comparative framework aims to uncover evolutionary mechanisms influencing the maintenance or erosion of genetic diversity, providing insights to inform conservation priorities for these and other vulnerable species.

## Materials and Methods

### Dataset

The reference genomes from three bird species endemic to Mauritius—the echo parakeet (Morales, Groombridge, et al. 2024), the pink pigeon (Morales, Van Oosterhout, et al. 2024), and the Mauritius kestrel (Morales, Norris, et al. 2024)—were recently sequenced and reported. To build a comparative dataset, 36 additional bird species were selected based on the availability of high-quality reference genomes (i.e., based on N50, average scaffold size, and total scaffold count) and ensuring a comprehensive representation across the avian phylogeny (Figure 1B). Metadata for all 39 species, including current census population sizes and IUCN conservation statuses, are compiled in Table S1.

### Mapping and variant calling

Raw reads used for assembling the reference genomes were downloaded from NCBI (see Table S1 for Assembly IDs) and aligned to the corresponding genomes. NGS short reads were mapped using BWA (v0.7.17) mem (Li and Durbin 2009) with default parameters. Read duplicates were marked with GATK (4.4.0.0) MarkDuplicates (DePristo et al. 2011). PacBio HiFi reads were mapped and sorted using pbmm2 v1.5.0 (https://github.com/PacificBiosciences/pbmm2) with the parameter “--preset HIFI”. GATK HaplotypeCaller was used to call variants for each alignment. Only SNPs were kept for further analyses.

### Coverage

After mapping the raw reads to each reference genome, we estimated the average genome-wide coverage per scaffold in each genome using MosDepth v0.3.3 (Pedersen and Quinlan 2018).

### Sex chromosome removal

Each reference genome was mapped to the chicken genome (assembly GRCg6a) with minimap2 v2.1 (Li 2018). Any scaffold mapped to chicken sex chromosomes for more than 20% was treated as potential regions from sex chromosomes and removed for further analyses. Any additional scaffolds annotated in the reference genomes as sex chromosomes were also removed.

### Heterozygosity

We estimated genome-wide heterozygosity using ANGSD (Korneliussen et al. 2014). We first obtained genotype likelihoods on scaffolds larger than 500 kb and only considered sites with a depth of coverage between ⅓ (-setMinDepth) and two times (-setMaxDepth) the average coverage for each sample. We assumed that the reference and ancestral states were the same. We applied the following parameters: -uniqueOnly 1 -remove_bads 1 - only_proper_pairs 1 -C 50 -baq 0 -minMapQ 30 -minQ 20 -setMinDepth $minDP - setMaxDepth $maxDP -doCounts 1 -GL 2 -doSaf 1. Next, we calculated the folded site frequency spectrum (SFS) with realSFS.

To compare different estimations of heterozygosity, we also estimated genome-wide heterozygosity directly from the VCF files with a custom pipeline. First, we divided scaffolds into sliding windows of size 100 kb (with a slide of 50 kb) using bedtools makewindows v2.30.0 (Quinlan and Hall 2010). Next, we obtained the total number of callable heterozygous and total genotypes per window using bcftools v1.20 (Danecek et al. 2021), tabix v1.14 (Li 2011), and vcfhetcount from vcflib (Garrison et al. 2022). Genotypes were considered callable if their read depth was between ⅓ and 2 times the average coverage per sample and had a minimum genotype quality (GQ) of 30 or reference genome quality (RGQ) of 10. Indels and windows with less than 50% of callable sites were excluded from the analysis.

### Runs of Homozygosity (ROH)

As some reference genomes were assembled with only Pacbio long reads, and no short-read data are available for the same individual, commonly used methods like ROHan (Renaud et al. 2019) could not be applied to identify ROH. To address this limitation, we developed a custom method for this analysis. Given the varying fragmentation levels of the assemblies (Table S1), for each species, we retained only scaffolds with a minimum size of 5 Mb. As a result, two species (Red-faced mousebird, *Urocolius indicus* and Wilson’s storm petrel, *Oceanites oceanicus*) were excluded, as neither had scaffolds larger than 5Mb. We identified ROH based on per-window heterozygosity estimates (see above). Using the R package bedtoolsr (Patwardhan et al. 2019), we concatenated windows with a heterozygosity lower than 5e^−4^ bp^−1^ (Figure S1), except for two genomes with exceptionally low average heterozygosity (the flightless cormorant, *Nannopterum harrisi*, and Mauritius kestrel), for which a lower threshold of 1e^−4^ bp^−1^ was used. Adjacent homozygous regions were merged if separated by a gap shorter than 100 kb, and only ROHs with a minimum size of 500 kb were retained. The inbreeding coefficient (F_ROH_) was calculated as the ratio of the total length of ROH segments to the total length of the analyzed genome (scaffolds > 5 Mb). We also estimated heterozygosity for the analyzed scaffolds per species, both including and excluding ROHs.

To validate our custom method, we compared our ROH estimates with those from ROHan for five species for which short-read data were available. ROHan was employed using a window size of 100Kb and a rohmu of 5e^−4^ to mimic the parameters used in the custom method.

### Genetic load

To compare genetic load across species, we used chicken CADD score annotations as a proxy for substitution deleteriousness at ultraconserved elements (UCEs) in our target species. We extracted UCE regions and corresponding flanking regions from each reference genome and from the chicken genome (GRCg6a), and performed a multi-species alignment for each UCE, using the recommended pipeline of Phyluce v1.7.3 (Faircloth 2016)see Figure S2 for detailed pipeline). Each genome was converted from fasta to 2bit format using UCSC FaToTwoBit (Casper et al. 2018). The UCE probe file was downloaded from https://github.com/faircloth-lab/uce-probe-sets/tree/master/uce-5k-probe-set, and used to extract and validate UCE regions per species. The CADD score file of the chicken genome was downloaded from https://osf.io/c97ez. For each species, CADD scores were lifted from the chicken genome for homozygous (relative to chicken) and heterozygous sites across UCE regions with customised scripts partially adapted from LoadLift (Speak et al. 2024). Heterozygous sites were extracted from VCF files and filtered using bcftools with the same parameters used for heterozygosity estimations (see above). Only heterozygous sites with one allele equal to the corresponding site in the chicken genome were kept for further analyses.

To quantify genetic load in each genome, we counted the number of heterozygous sites and homozygous substitutions with CADD scores ⩾ 20 in UCE regions, representing the top 1% most deleterious sites in the chicken genome. To control for the the evolutionary distance to the chicken among species we rescaled the counts of sites with CADD scores ⩾ 20 with the counts of substitutions with CADD scores < 3, the latter representing nearly-neutral sites. To account for potential lineage-specific adaptive substitutions that have occurred since the divergence with chicken, homozygous substitutions that were shared by 20 or more species were excluded, as these are more likely to reflect long-term adaptive changes rather than harmful mutations.To further confirm that we were correctly retaining sites that have not changed since the divergence with chicken, we compared three species to their closest reconstructed ancestral nodes in the phylogenetic tree from Stiller et al. (2024). We extracted the reconstructed full sequences from the closest ancestral nodes and cut them into 200-bp short sequences with 20-bp step sliding windows. The reads were mapped to the chicken genome with BWA. For each species, we confirmed that homozygous substitutions had at least one ancestral read mapped to the chicken genome, and the ancestral state matched the chicken sequence. Homozygous substitutions with CADD scores above 20 are concentrated at the terminal branches of the phylogenetic tree of birds (Figure S3), indicating that they are more likely to be deleterious substitutions than lineage-specific adaptations. The rescaled relative ratio of heterozygous sites was used as a proxy of masked load, which we refer as “heterozygous load” in the following sections. The relative ratio of homozygous sites, being referred to as “homozygous load”, was used as a proxy realized load.

### Demographic history

We inferred historical fluctuations of effective population size (Ne) for each species using PSMC (Li and Durbin 2011) with the parameters “-N30 -t5 -r5 -p 1+1+1+1+30*2+4+6+10” (Hilgers et al. 2025) and estimated the harmonic Ne mean from 10 kya to 100 kya for further analyses.

### Statistical and phylogenetic comparative analyses

To explore the relationships among genome-wide heterozygosity, F_ROH_, Ne, and genetic load across species while accounting for phylogenetic signals, we reconstructed the phylogenetic relationships among the studied species and incorporated this inference into two statistical frameworks. The phylogeny was reconstructed from a subset of the UCE dataset described above (see *Genetic Load* section). Using the software AMAS (Borowiec 2016), we selected and concatenated 1,526 UCE sequences, retaining only loci with ≤2% missing data and >30% parsimony-informative sites. We used IQ-TREE (Minh et al. 2020) to infer the maximum-likelihood tree using the edge-linked partition model (Chernomor et al. 2016) and constraining the topology to reflect higher-order taxonomic relationships following Stiller et al. (2024). Using this inferred phylogenetic tree, we assessed different univariate models with a phylogenetic generalised least squares (PGLS) approach (Freckleton et al. 2002). These analyses were conducted using the R packages *ape* (Paradis et al. 2004; Paradis and Schliep 2019), *caper* (Orme et al. 2023) and *nlme* (Pinheiro et al. 2014). When PGLS indicated a non-zero lambda (λ) value, suggesting a significant phylogenetic signal, we further examined these effects using a Bayesian phylogenetic generalised linear mixed model (pGLMM) (Hadfield and Nakagawa 2010). In this framework, the phylogenetic relationship among species was modelled as a random effect. pGLMM analyses were performed using the R packages *ape* and *MCMCglmm* (Hadfield 2010).

### Genomic synteny

We inferred multi-genome synteny for chromosome-level reference genomes for pigeons (three species in Columbidae), parrots (five species in Psittaciformes), and falcons (six species in *Falco*) separately using ntSynt v1.0.2 (Coombe et al. 2024) with the divergence range (-d) set to 10. Synteny results were visualised using scripts from ntSynt based on the R package *gggnomes* (Hackl et al. 2024).

### Identification of Transposable Elements

To annotate repetitive elements (RE) in the genomes of the pink pigeon, Mauritius kestrel, and Mauritius parakeet, we produced *de novo* libraries of RE for each species using RepeatModeler2 (Flynn et al. 2020). We combined the *de novo* libraries with previously published manually curated libraries of RE from the Collared flycatcher (*Ficedula albicollis*) and Blue-capped cordon-bleu (*Uraeginthus cyanocephalus*) from Storer et al. (2021), and from the Emu (*Dromaius novaehollandiae*), Anna’s hummingbird (*Calypte anna*), and Kākāpō (*Strigops habroptilus*) from Peona et al. (2021). Using the resulting custom libraries, we annotated the RE from the genomes using RepeatMasker version 4.0.8 (Smit et al. 2015). We repeated this process for three, five and four additional species of Columbiformes, Falconiformes and Psittaciformes, respectively, to enable comparisons of proportions of RE across species.

## Results

### Genome-wide diversity and inbreeding

The sequencing depth across the compiled dataset ranged between 15x and 96x (mean = 46, SD = 19). We estimated genome-wide heterozygosity with both genotype likelihoods in ANGSD and by SNP-calling, resulting in very similar estimates (adjusted R^2^ = 0.78; Table S1, S2, Figure S4). We used heterozygosity estimates from ANGSD for all subsequent analyses, except for the estimation of ROHs (see Material and Methods). Neither heterozygosity or F_ROH_ estimates showed significant correlation with the quality of the genomes (e.g., N50) or depth (Figure S5). Our in-house method produced consistent ROH results to those from ROHan (Figure S6), validating our approach.

Genome-wide heterozygosity showed a strong negative correlation with inbreeding coefficient F_ROH_ (Figure 1C; PGLS: λ = 0, R^2^ = 0.46, F_1-28_ = 26.56, *p* < 0.001). As expected, samples with domestic or pet origins (Table S1) showed higher F_ROH_ compared to wild samples with similar levels of heterozygosity. Thus, we excluded the former samples from further analyses. IUCN Red List status was not a good predictor of genome-wide heterozygosity nor of F_ROH_ (Figure S7) or genetic load (Figure S8). The log-transformed Ne/Nc ratio (Log(Ne)/Log(Nc)) was significantly lower (Wilcoxon two-sample test *p* < 0.001) in non-threatened species (Least Concern, mean = 0.91, SD = 0.23) compared to threatened species (remaining IUCN status categories, mean = 1.49), yet more varied (SD = 0.52). When numbers are assigned to the status categories (0 to Critically Endangered, 1 to Endangered, …, 4 to Least Concern), Ne/Nc ratios had a linear correlation with IUCN status (Figure 1D, GLM: R^2^ = 0.43, F_1-24_ = 12.49, p < 0.001). The Ne value (estimated with PSMC) represents an estimate of ancestral population size deep-in-time, while the Nc value represents current census population size. Elevated Ne/Nc ratios are indicative of recent population declines, with higher values reflecting more abrupt changes. This result illustrates the discrepancy between IUCN Red List conservation status relying only on demographic estimates and the shallow correlation to the genetic diversity estimates.

We found a significant positive correlation between historical Ne and genome-wide heterozygosity (Figure 2A; PGLS: λ = 0, R^2^ = 0.47, F_1-22_ = 19.65, *p* = 2.1e^−4^), highlighting the predominant effect of long-term demographic history on genetic diversity. In contrast, F_ROH_ showed no correlation with historical Ne (Figure 2D; PGLS: λ = 0, R^2^ = 0.11, F_1-22_ = 2.82, *p* = 0.11). This lack of correlation is not surprising, given that our analysis focused only on long ROHs (> 500 Kb), which reflect population history within tens to hundreds of generations ago. Species with higher genetic diversity or a lower inbreeding coefficient tended to be burdened by a higher heterozygous load, measured as the corrected ratio of heterozygous sites with a CADD score above 20 (Figure 2B; PGLS: λ = 0, R^2^ = 0.61, F_1-30_ = 45.99, *p* = 1.6e^−7^, and Figure 2E; PGLS: λ = 0, R^2^ = 0.91, F_1-28_ = 6.74, *p* = 0.015). In contrast, homozygous load, measured as the corrected ratio of homozygous substitutions with a CADD score above 20, showed a statistically significant but weak association with heterozygosity (Figure 2C; PGLS: λ = 0.81, R^2^ = 0.16, F_1-30_ = 5.61, *p* = 0.0245), and no association with F_ROH_ (Figure 2F; PGLS: λ = 0.86, R^2^ = 0.02, F_1-28_ = 0.61, *p* = 0.4424). However, given the detected phylogenetic signal in the homozygous load comparisons (λ ≈ 0.8), we further explored these relationships using a PGLMM approach. The results supported the previous described associations and revealed that phylogenetic relatedness does not play a predominant role in homozygous load variation (see Table S3).

**Figure 2.**
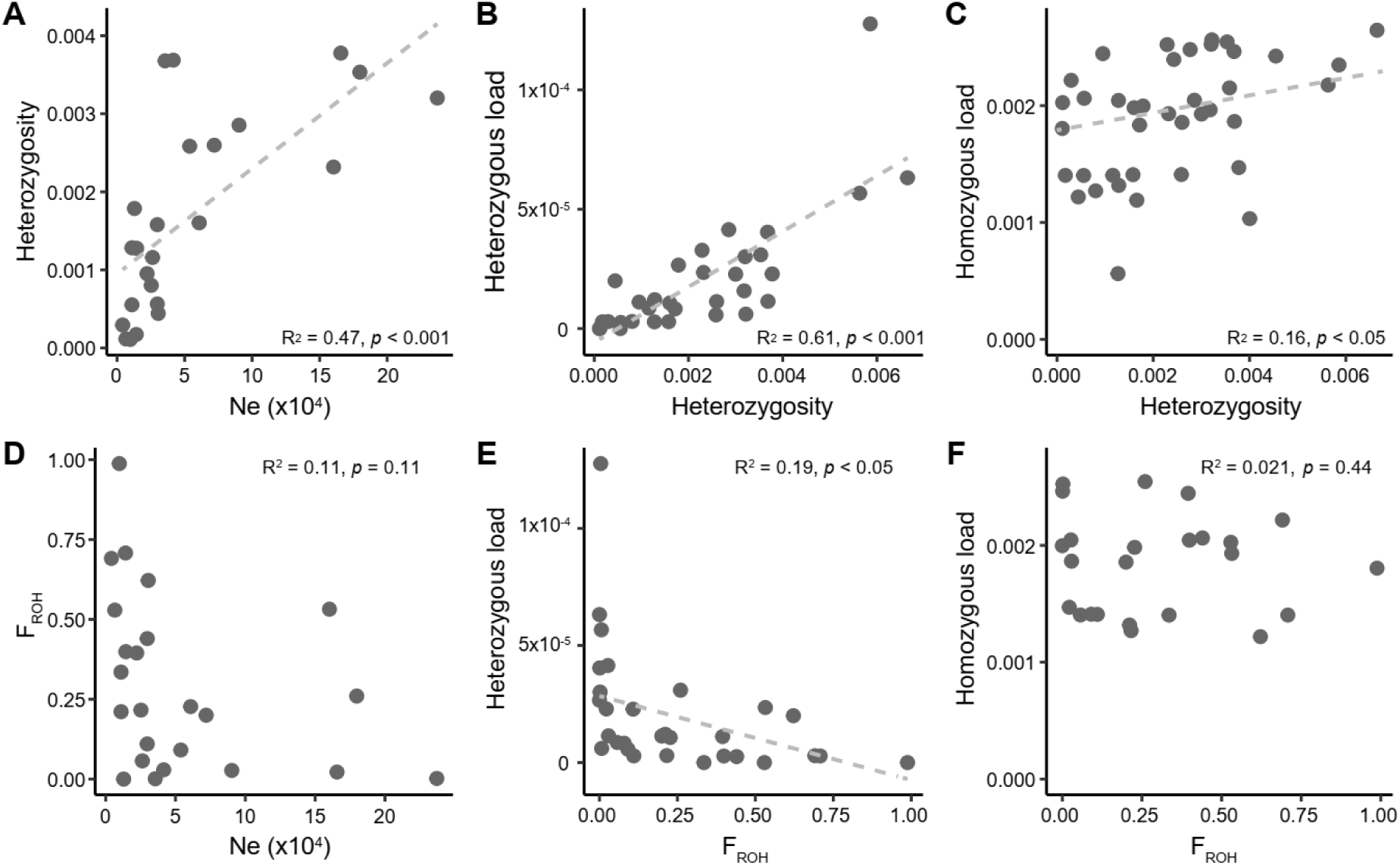
Comparison between genetic diversity metrics, genetic load, and effective population sizes. The dashed line represents the linear correlation when *p*<0.05. **A** Correlation between genome-wide heterozygosity and effective population size (Ne), with Ne estimated as the harmonic mean of PSMC values between 10 kya and 100 kya. **B** Correlation between heterozygous load and genome-wide heterozygosity. Heterozygous load is defined as the ratio of heterozygous substitutions with CADD scores above 20 to homozygous substitutions with CADD scores below three. **C** Correlation between putatively homozygous load and genome-wide heterozygosity. Homozygous load is defined as the ratio of counts of filtered homozygous substitutions with CADD scores above 20 and the number of filtered homozygous substitutions with CADD scores below three. **D** Correlation between inbreeding coefficient (based on runs of homozygosity; F_ROH_) and Ne. **E** Correlation between heterozygous load and inbreeding coefficient. **F** Correlation between putatively homozygous load and inbreeding coefficient.

### Inbreeding, genetic diversity and load across taxonomic groups

Despite similar distribution ranges and histories of population decline in the past decades (Figure 1A), the three Mauritius species showed contrasting levels of genetic heterozygosity, inbreeding coefficients (Figure 1C, 3AB), and homozygous load (Figure 3C). The Mauritius kestrel had the lowest heterozygosity of all samples (8.33 × 10⁻⁵ het x bp^−1^) and the second highest F_ROH_ (0.71) among the wild species included in this study, with 50% of its genome in very long ROHs (>10 Mb), as evidence of sustained recent inbreeding after recovering demographically from a bottleneck of only four individuals. The echo parakeet had the second lowest heterozygosity (8.07×10⁻⁴ het x bp^−1^) among parrots, closely following another extremely bottlenecked species, the critically endangered Kākāpō (*Strigops habroptila*). However, the echo parakeet’s F_ROH_, while high (0.40), is lower than that of the Kākāpō (0.69), with 24.7% of their genome in ROHs longer than 10 Mb, evidence of their extreme bottleneck of ∼12 individuals. The pink pigeon exhibits a heterozygosity of 2.38 × 10⁻³ het x bp^−1^, the lowest among the analysed pigeons, but higher than that of more than half of the species included in the study and nearly 30 times greater than that of the Mauritius kestrel. Additionally, the pink pigeon showed an F_ROH_ of 0.26, with 12.3% of its genome in ROHs of a length longer than 10 Mb (Figure S9).

**Figure 3.**
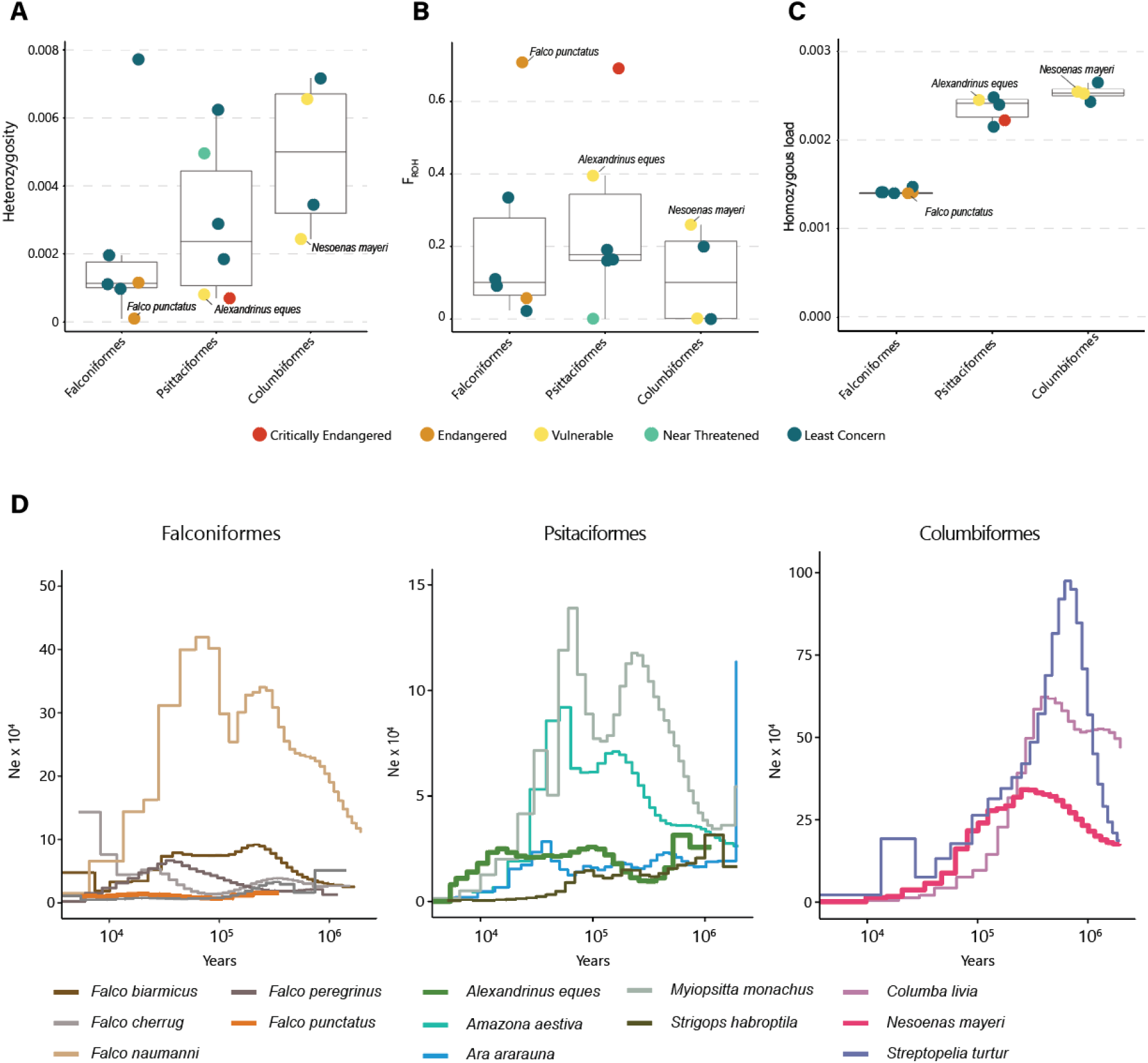
Genetic diversity, genetic load, and demographic history of Falconiformes, Psitaciformes, and Columbiformes. **A** Genome-wide heterozygosity in het x bp^−1^, **B** inbreeding coefficient, and **C** homozygous load distribution across the orders of the three target species. The inbreeding coefficient (F_ROH_) was estimated using runs of homozygosity, and homozygous load was based on the ratio of homozygous substitutions with CADD scores above 20 to those with CADD scores below 3. **D** Demographic histories are shown as variation in Ne (effective population size) inferred with PSMC. Thick lines refer to Mauritius species. Only species with chromosomal-level assemblies were included.

The differences in genetic diversity of Mauritius species were associated with the differences between their taxonomic groups (Figure 3). Falcons exhibit the lowest heterozygosity and homozygous load among the three taxonomic groups. Likewise, falcons carry 23.7% less homozygous load than the parrots and 29.7% less than the pigeons. Within their respective taxonomic groups, the three Mauritius species showed the lowest genome-wide heterozygosity (Figure 3A) and the highest F_ROH_ (Figure 3B). Genetic diversity estimates carry the signal of ancestral population size, as the three Mauritius species had relatively low population sizes within their respective taxonomic groups (Figure 3D). However, the pink pigeon had a larger historical population than most studied species, including falcons and parrots, which is reflected in its higher heterozygosity compared to the average levels in the other taxonomic groups (Figure 3A). This reveals the importance of considering genetic diversity within the context of a species’ long-term evolutionary history and taxonomic group. Note that all species that we have included from Falconiformes are from the same genus, *Falco*, with a divergence time of 12 million years (Kumar et al. 2022), which could explain the small deviations for heterozygosity and homozygous load, whereas the divergence time of the study Columbiformes and Psittaciformes species was roughly 16 and 55.6 milion years, respectively (Figure 4).

**Figure 4.**
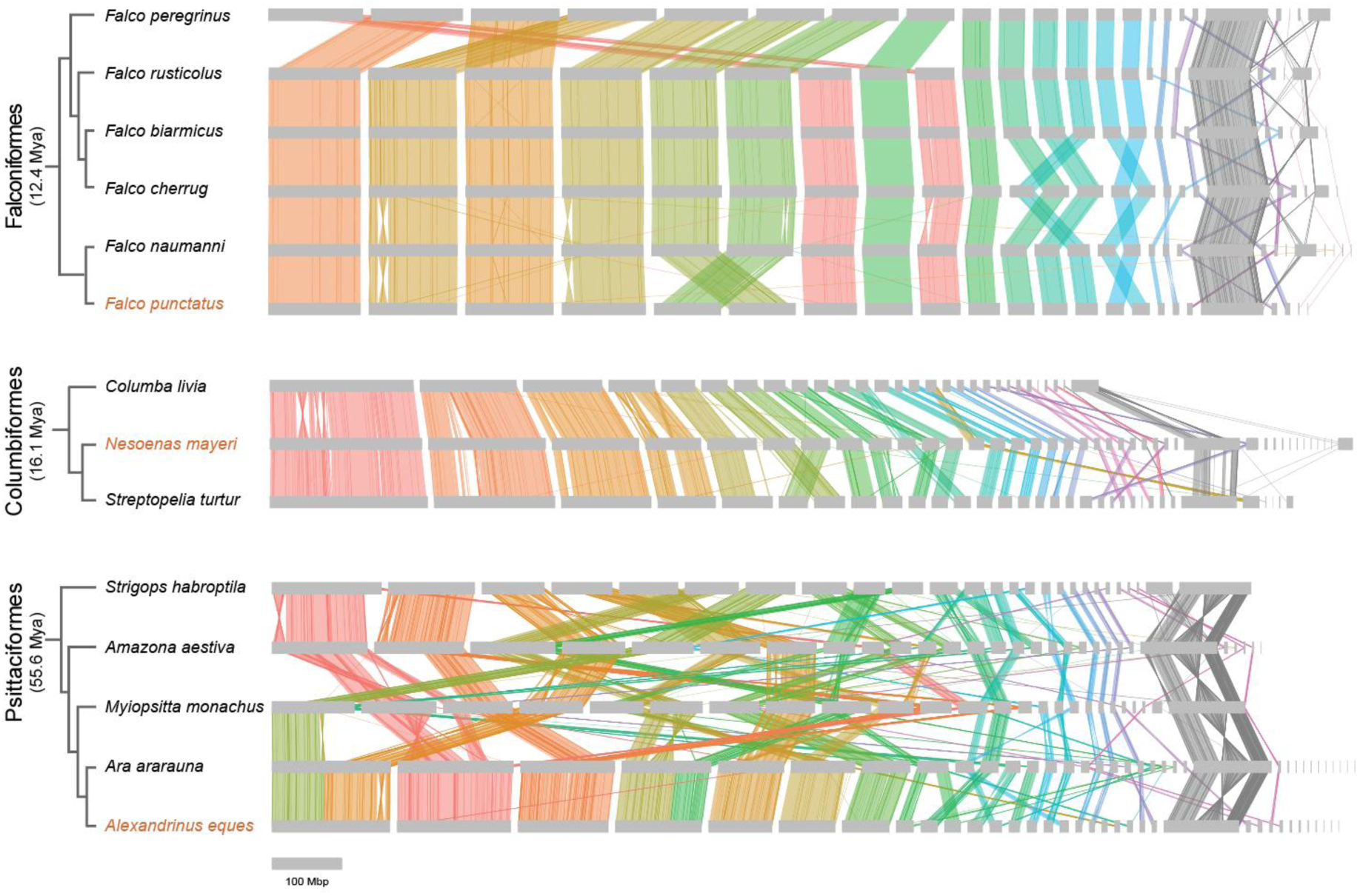
Pattern of chromosome synteny within Falconiformes, Columbiformes and Psittaciformes. For each species, chromosomes (grey horizontal bars) are ordered according to their respective genome assemblies. Syntenic relationships (i.e., syntenic blocks identified by ntSynt) between macro- and medium-sized chromosomes are marked with differently coloured vertical lines, while synteny between sex chromosomes and micro-chromosomes is marked with grey vertical lines. On the left, the phylogenetic relationship among species is provided (cladogram; branch lengths not scaled to time), with the time to the most recent common ancestor (millions of years ago) indicated for each group of species. The names of the three Mauritius species are in orange.

**Figure 5.**
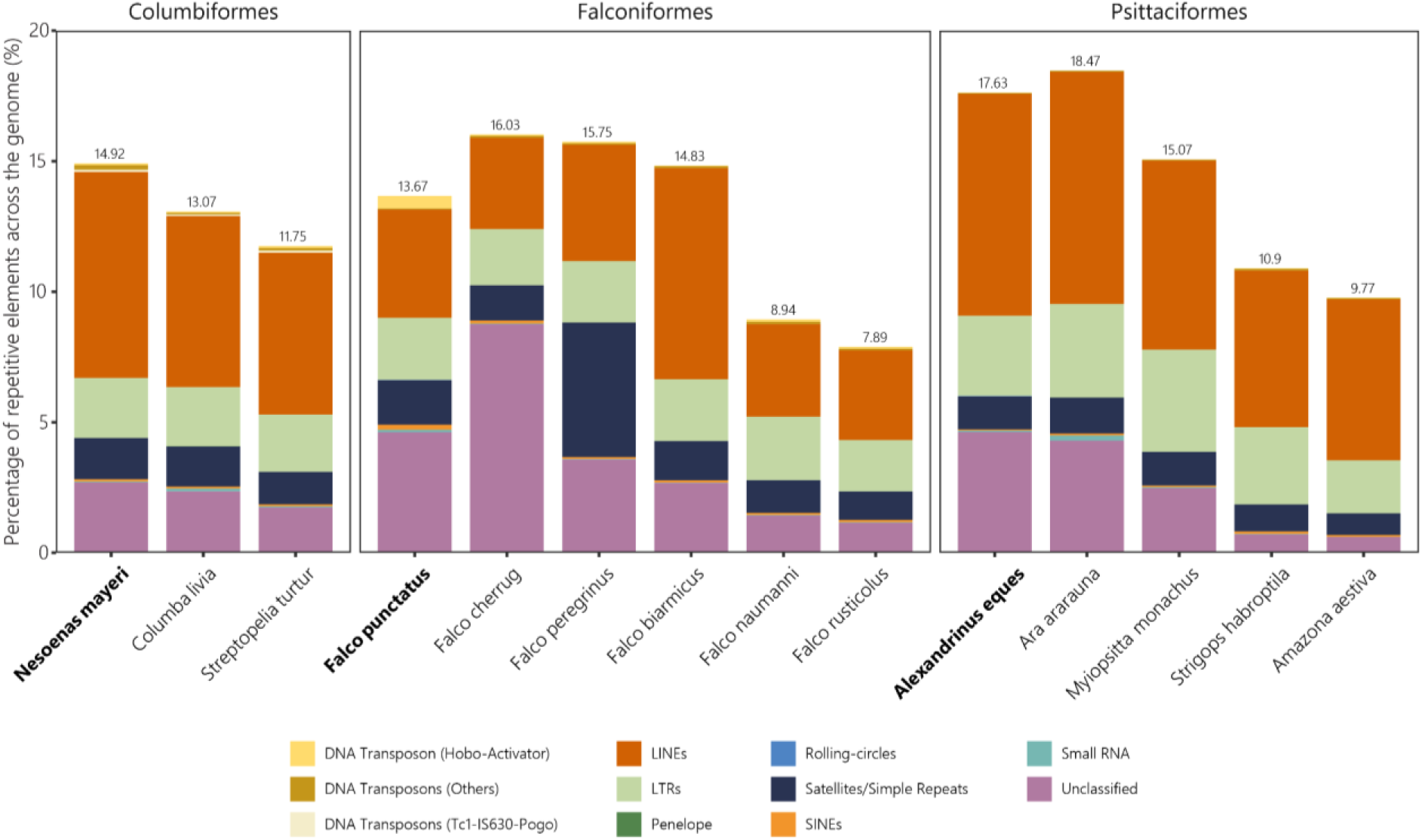
Annotated repeated elements (RE) in Columbiformes, Falconiformes and Psittaciformes. The proportion of different classes of RE in the genome (different colours) and the total proportion of RE (value on top of each bar) are shown for each species.

### Genome synteny and repetitive elements

The three taxonomic groups differed in degree of genome conservation, with Falconiformes and Columbidae having relatively stable genomes, while Psittaciformes showed highly complex rearrangements between most chromosomes and species (Figure 4). However, also within Falconiformes and Columbidae, individual chromosomes showed evidence of rearrangement. For example, Chr1 of the peregrine falcon (*Falco peregrinus*) was homologous to Chr7 and Chr9 of the other *Falco* species (indicating a fusion or a fission), and the lesser kestrel (*Falco naumanni*) showed evidence of intra-chromosomal inversions on both Chr2 and Chr4. In general, closely related species are expected to show maintained synteny, and in line with this reasoning, the three Mauritius species showed well-maintained synteny to the closest related species included in the comparison (Figure 4).

### Repetitive Element Annotation

The average percentage of repetitive elements (RE) was 13.25% for the Columbiformes (14.92% for the pink pigeon, highest among the studied pigeons), 12.85% for the Falconiformes (13.67% for the Mauritius kestrel), and 14.37% for the Psittaciformes (17.63% for the Mauritius parakeet). These results align with previous studies suggesting a lower proportion of RE in avian genomes compared to those in mammals, with most avian species presenting around 15-20% of RE (Hughes and Piontkivska 2005). All Columbiformes presented the transposon Tc1.IS630, which was not present in Psittaciformes and Falconiformes. This transposon element has been involved in structural rearrangement and has a gene regulatory function (Shen et al. 2021; Wang et al. 2021).

In the falcons, we found a sharp difference of almost 5% in RE between the sequenced species using PacBio Hi-Fi (N = 13) and Illumina short-read sequencing (N = 1, *Falco cherrug*) techniques (Table S1). It has been shown that short-read sequences are not ideal for annotating RE due to their inherent fragmentation (Mann et al. 2024). However, the three Mauritius species presented in this study have been long-read sequenced. The Mauritius kestrel presented the highest percentage (0.48%) of the transposon hobo activator compared to all species analyzed. The hobo transposon has been associated with developmental regulatory genes, suggesting a role in the evolution of developmental gene networks (Deprá et al. 2009). It can induce transposition through the “cut and paste” mechanism, which can impact developmental processes (Kim et al. 2011).

In the parrots, the Mauritius parakeet presented the second highest percentage of RE (17.63%) compared to the rest of the Psittaciformes studied, highlighting potential lineage-specific retention or amplification of repetitive elements. In contrast, the blue-fronted amazon (*Amazona aestiva*) and kākāpō (*Strigops habroptila*) exhibited markedly lower RE percentages relative to other species. Given the functional implications of RE in genome organization and regulatory evolution, these differences may reflect distinct evolutionary pressures shaping the genomic architecture of these taxa.

## Discussion

By analysing 39 genomes from a diverse range of avian species, we demonstrate that deep demographic history exerts a lasting influence on genetic diversity and, consequently, on masked genetic load. Species from different taxonomic groups exhibit different levels of heterozygosity, and while their demographic history can account for some of these differences, this also suggests that there might be taxonomic-specific differences in life-history traits that exert an influence on how genetic diversity and inbreeding respond to demographic decline and recovery. Overall, our results highlight the importance of accounting for demographic history and taxonomy in interspecies comparisons of genetic metrics. While we identified some challenges for standardizing genomic metrics comparisons across diverse taxonomic groups, our study demonstrates that a comparative genomics framework offers powerful insights for addressing key questions in biodiversity conservation.

### Understanding conservation genetics with comparative methods

Leveraging the rapidly expanding availability of high-quality avian genomes (Feng et al. 2020; Stiller et al. 2024), we are now able to employ comparative phylogenetic methods to gain deeper insights into genetic traits that play important roles in conservation biology (Supple and Shapiro 2018; Wright et al. 2020). We observed a negative correlation between genetic diversity and F_ROH_ (Figure 1C), consistent with previous studies (Brüniche-Olsen et al. 2018; Grossen et al. 2020), showing a higher risk of genetic drift and inbreeding in small populations. Reference genomes sequenced from domestic or pet samples had elevated F_ROH_, indicating the importance of checking the resources of the samples when using public genomic data (Figure S9).

Data on modern population sizes are undoubtedly crucial in conservation assessments (Shaffer 1981; Lande 1988; Willi et al. 2006; Frankham et al. 2014). The rate of decline is one of the major factors considered by the IUCN Red List rating (Frankham et al. 2014; McNeely et al. 1990). The Ne/Nc (with modern Ne) ratio reflects the balance between genetic diversity and current population size (Frankham 1995; Kalinowski and Waples 2002), with an increased Ne/Nc ratio suggesting a rapid population decline that has not yet resulted in significant genetic diversity loss measured as genome-wide heterozygosity. This highlights the prevalent time-lag between population decline and genetic diversity loss (Gargiulo et al. 2024) resulting from the drift debt (Gilroy et al. 2017; Dussex et al. 2023; Pinto et al. 2024; Liu et al. 2025), serving as an early warning sign of an imminent population collapse (Amos and Balmford 2001; Wilder et al. 2023). Our findings indicate that the Ne/Nc ratio, even with historical Ne, serves as an indicator of a species’ conservation status (Figure 1D), showing the importance of understanding the long-term demographic history of the species in conservation.

Genetic diversity has been considered a classical indicator for population resilience and risk of extinction (Breed et al. 2019; DeWoody et al. 2021; Teixeira and Huber 2021; Jeon et al. 2024). We showed that genome-wide heterozygosity, as a measurement of genetic diversity, strongly correlates with historical population size (Figure 2A) spanning 10,000 to 100,000 years ago, with the former corresponding to at least 500 to 5,000 generations in birds given a generation time of roughly 2-20 years (Bird et al. 2020). This indicates a species’ “genetic vulnerability” to future challenges is associated with its long-term evolutionary history, even prior to the accelerated environmental changes induced by human activity (Tan et al. 2023). Thus, monitoring both population demography and environmental threats could be important to the conservation of species and populations with low genetic diversity, even if the population sizes remain stable, as they are initially more likely to be vulnerable to environmental changes (Ellstrand and Elam 1993; van der Valk, de Manuel, et al. 2019; Brüniche-Olsen et al. 2021; Liu et al. 2025; Willi et al. 2006). F_ROH_, on the other hand, did not show a strong correlation with the historical population size, as long runs of homozygosity reflect recent demographic history within tens of generations (Ceballos et al. 2018), highlighting the importance of combining different genomic metrics to evaluate the genetic health of a species and their likely short- and long-term risks.

As whole-genome data become more accessible, the integration of genomic-derived metrics (e.g., demographic reconstructions, heterozygosity, F_ROH_, Ne/Nc) with demographic, ecological and environmental metrics can substantially improve conservation assessments and planning. Combining genomic insights with ecological data will enable more precise predictions of population collapse and help prioritise efforts for species most at risk.

### A comparative perspective on genetic load

Population decline often leads to the expression of masked genetic load, driven by genetic drift and reduced purging (van der Valk, de Manuel, et al. 2019; Dussex et al. 2023). Here, we found a strong correlation between heterozygous deleterious sites, as a proxy for masked load, and genome-wide heterozygosity (Figure 2B). This suggests that species with higher genetic diversity face a different type of threat under demographic and environmental changes than those with lower genetic diversity. With habitat loss predicted to intensify in the near future, resulting in accelerated population declines and loss of genetic diversity (Exposito-Alonso et al. 2022), species with currently higher diversity may face rapid exposure of these deleterious mutations before effective purging can occur, increasing the risk of fitness reduction and jeopardising population viability (van Oosterhout et al. 2022).

Conversely, species with lower genetic diversity exhibit reduced homozygous load (Figure 2C). Despite that one sample cannot reflect the whole picture of a species, this pattern across a wide range of species likely reflects long-term purging of deleterious alleles in populations with small Ne. Although we cannot rule out lineage-specific adaptive substitutions when measuring homozygous load, focusing on sites with CADD > 20 within the most conserved regions provides a reasonable control (Rentzsch et al. 2019; Fontsere, Speak, Caven, Rodriguez, et al. 2024; Speak et al. 2024) (Figure S3). However, such purging does not necessarily translate to improved fitness (Kardos et al. 2021). If this interpretation is correct, it implies that the signal we measured primarily captures mildly deleterious sites, as strongly deleterious alleles are typically purged quickly (Robinson et al. 2016; Dussex et al. 2021; Dussex et al. 2023; Fontsere et al. 2024), and our estimation included substitutions reflecting the accumulation of putatively deleterious homozygous alleles during a long evolutionary process, rather than a potential negative fitness effect. This also suggests that genetic load analyses tend to detect mainly mildly deleterious alleles as a proxy of the realized load (Grossen et al. 2020; Dussex et al. 2023; Kardos et al. 2023; Wang et al. 2023). Addressing this challenge requires improving ancestral state inference, which currently relies on species that diverged tens of thousands of generations ago.

Moreover, of key importance is advancing our understanding of the fitness effects of putatively deleterious alleles to better integrate genetic load estimation into conservation strategies (Kardos et al. 2021; Dussex et al. 2023). This could involve incorporating direct fitness data, leveraging temporal genomic data to trace load dynamics, and refining methods for detecting strongly deleterious alleles (Bosse et al. 2019; van der Valk, de Manuel, et al. 2019; Bertorelle et al. 2022; Kyriazis et al. 2023). Understanding the interplay between genetic load, structural variation, and demographic history will be crucial for predicting species’ resilience to environmental change and guiding effective management.

### Comparative genomics reveals patterns across taxonomic groups

The three Mauritius species exhibit low genetic diversity and high F_ROH_ within their species groups, as expected from their recent histories of population collapse. However, substantial genetic differences between their taxonomic groups highlight the interplay of evolutionary history, demographic processes, life-history traits and genomic architecture. For instance, the pink pigeon exhibits higher heterozygosity than most *Falco* species (Figure 3A), whereas *Falco* species show significantly lower homozygous load compared to parrots and pigeons (Figure 3C). These patterns underlie the effect of long-term effective population sizes (Figure 3D). On the other hand, F_ROH_ values based on long ROHs have similar values across taxonomic groups (Figure 3B), as these do not reflect the effect of long-term demographic history.

Life-history traits may help further explain these patterns. Parrots, such as the Mauritius parakeet, have long generation times, low reproductive rates, and high parental investment (Jones and Swinnerton 1997; Jones 2010; Jones et al. 2013), making them particularly vulnerable to genomic erosion. These traits slow the recovery of genetic diversity after bottlenecks and exacerbate the accumulation of homozygous load. In contrast, pigeons, such as the pink pigeon, have shorter generation times and higher reproductive rates (Jones 2013), facilitating faster recovery and preserving higher genetic diversity despite similar population collapses. Falcons, including the Mauritius kestrel, exhibit intermediate traits, with low reproductive rates but shorter generation times and the ability to disperse to new environments (Jones et al. 1995; Cartwright et al. 2014; Nicoll et al. 2021), which can limit genetic drift and inbreeding but may not fully mitigate the effects of historically small population sizes. However, the connection between life-history traits and genetic traits remains to be better studied (Germain et al. 2023).

Distinct patterns in genomic architecture further illustrate the importance of considering phylogenetic context in conservation genetics. Falcons exhibit reduced levels of Long Interspersed Nuclear Elements (LINEs), while parrots have higher proportions of Long Terminal Repeats (LTRs), suggesting taxonomic variation in transposable element composition (Kapusta and Suh 2017; Benham et al. 2024). These differences may influence key genomic processes, including gene regulation and alternative splicing, potentially shaping species’ adaptive capacities (Lin et al. 2009; Schmitz and Brosius 2011; Chénais et al. 2012; Casacuberta and González 2013). Similarly, parrots are known to experience more frequent chromosome reshuffling (Z. Huang et al. 2022), a pattern confirmed in our study (Figure 4). Chromosome rearrangements, such as inversions, may impact the evolution of genetic load by reducing recombination and preserving deleterious alleles (Jay et al. 2021; K. Huang et al. 2022). Beyond traditional metrics of genetic diversity, chromosome rearrangements and transposable elements may become more important in our understanding of genomic erosion, although their links to conservation outcomes remain underexplored. Future studies incorporating high-resolution genomic analyses across diverse taxa could help clarify the role of structural variation in shaping species’ evolutionary trajectories and their responses to ongoing environmental challenges (Brüniche-Olsen et al. 2021).

Our findings illustrate the importance of accounting for taxonomy, genomic architecture, and life-history traits when comparing species of conservation concern to closely related taxa (e.g., Robinson et al. 2018, 2019; Grossen et al. 2020). While comparative methods provide valuable insights, future research could investigate how life-history traits and genomic features, such as transposable elements and chromosome rearrangements, interact with genomic erosion and demographic change.

### Future directions to benefit conservation genomics

Our study underscores the value of expanding the availability of reference genomes to enhance the utility of genomic resources for conservation biology (Grueber 2015; Supple and Shapiro 2018). Given the substantial genetic differences observed between groups, it is advantageous, even in the absence of a species-specific reference genome, to identify a closely related reference genome to achieve better accuracy for certain analyses and inferences (Prasad et al. 2022). However, it is important to recognise that population genetic data are crucial, as a single individual cannot fully represent the genetic diversity of an entire species. Intra-species genetic variation and structure should be considered (Gutiérrez-Espeleta et al. 2000; Bowen et al. 2005; Turchetto et al. 2016), and relying on data from one or a few individuals may inadvertently capture outliers (Figure S9). Incorporating population-level data also allows for more robust estimates of realised genetic load by leveraging site frequency distributions (Grossen et al. 2020; Bertorelle et al. 2022).

Our results also emphasise the importance of integrating demographic history at multiple temporal scales to correctly interpret genetic diversity trends. Historical genetic data plays a critical role in accurately assessing trends in population size and genetic diversity (van der Valk et al. 2019; Femerling et al. 2023; Cavill et al. 2024; Dehasque et al. 2024; Silver et al. 2024; Fontsere et al. 2024). Such data are invaluable for identifying rapid declines in population size and genetic health, which may pose significant conservation risks of genetic erosion (Díez-del-Molino et al. 2018).

Comparative genomics offers a powerful framework for understanding how evolutionary history, demographic processes, and life-history traits shape genetic diversity across species (Bertorelle et al. 2022; van Oosterhout et al. 2022). By identifying commonalities and differences among taxonomic groups, this approach can inform targeted strategies, such as selecting species for genetic rescue or identifying populations most vulnerable to environmental change. Future research should explore how genomic features, such as genetic load, genome synteny and transposable elements, interact with population collapse and environmental change to influence species’ resilience (Díez-del-Molino et al. 2018; van Oosterhout et al. 2022; Germain et al. 2023). Moreover, integrating genomic data with ecological models and leveraging emerging tools like AI can provide a more holistic understanding of biodiversity conservation (van Oosterhout 2024).

## Acknowledgments

We are grateful to Anna Brüniche-Olsen and Roberto Biello for providing comments on an early draft version of the manuscript. This work was supported by the European Research Council (101078303); and the Swedish Research Council for Sustainable Development (2022-00536). Further support was obtained from the Royal Society International Collaboration Awards 2020 (ICA/R1/201194), the Earth and Life Systems Alliance (ELSA), the Swedish Research Council (621-4996), the Erik Philp-Sörensen’s foundation, Science for Life Laboratory (SciLifeLab), and Biodiversity and Ecosystem services in a Changing Climate (BECC). Views and opinions expressed are however those of the authors only and do not necessarily reflect those of the European Union or the European Research Council. Neither the European Union nor the granting authority can be held responsible for them.

## Data Availability

The scripts used in this study are available on GitHub: PachecoMC/CompConGen.

