## Supplemental Figures for "Genomic Erosion Through the Lens of Comparative Genomics"

* These authors contributed equally.

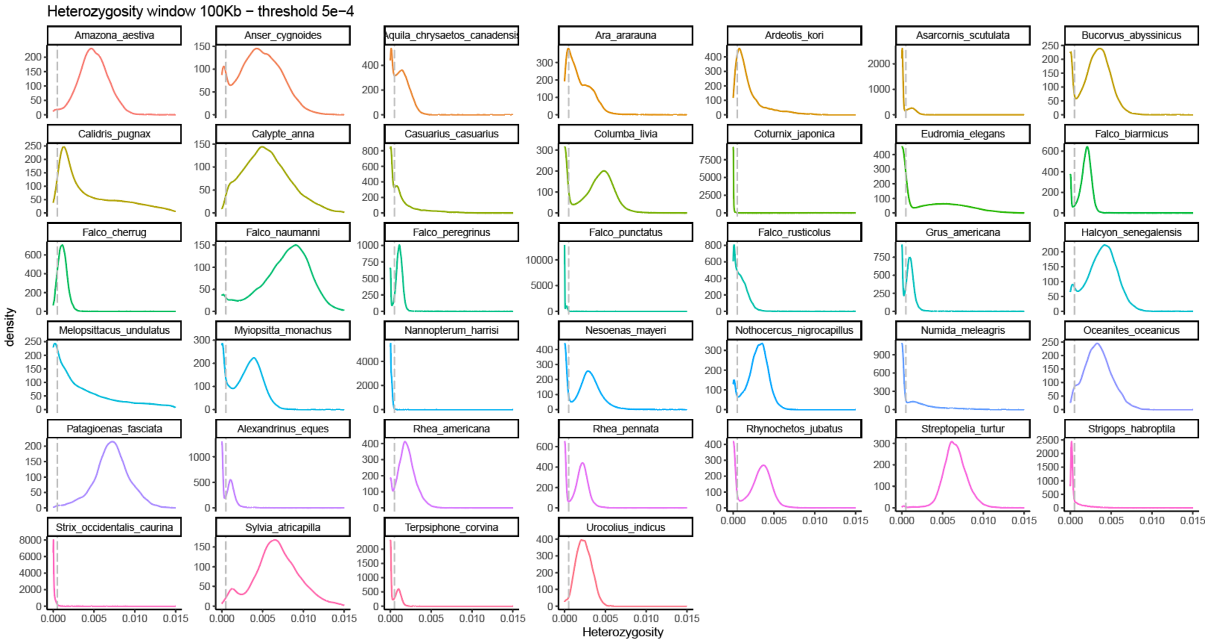


**Figure S1**. Distribution of heterozygosity in sliding windows of 100kb (50kb of slide). Vertical dashed gray line is set at 5e-4.

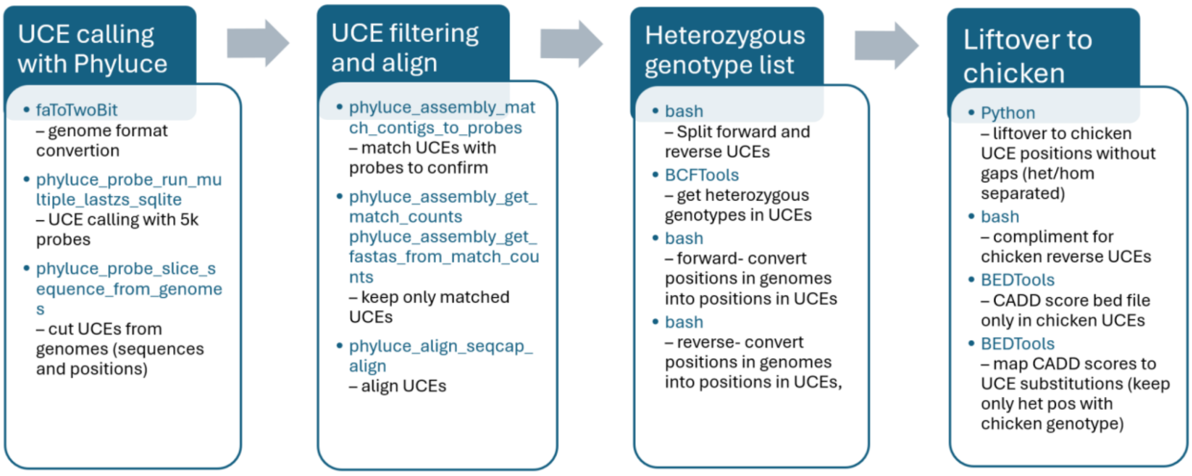


**Figure S2**. Detailed pipeline for calling UCE regions and CADD score liftover from chicken.

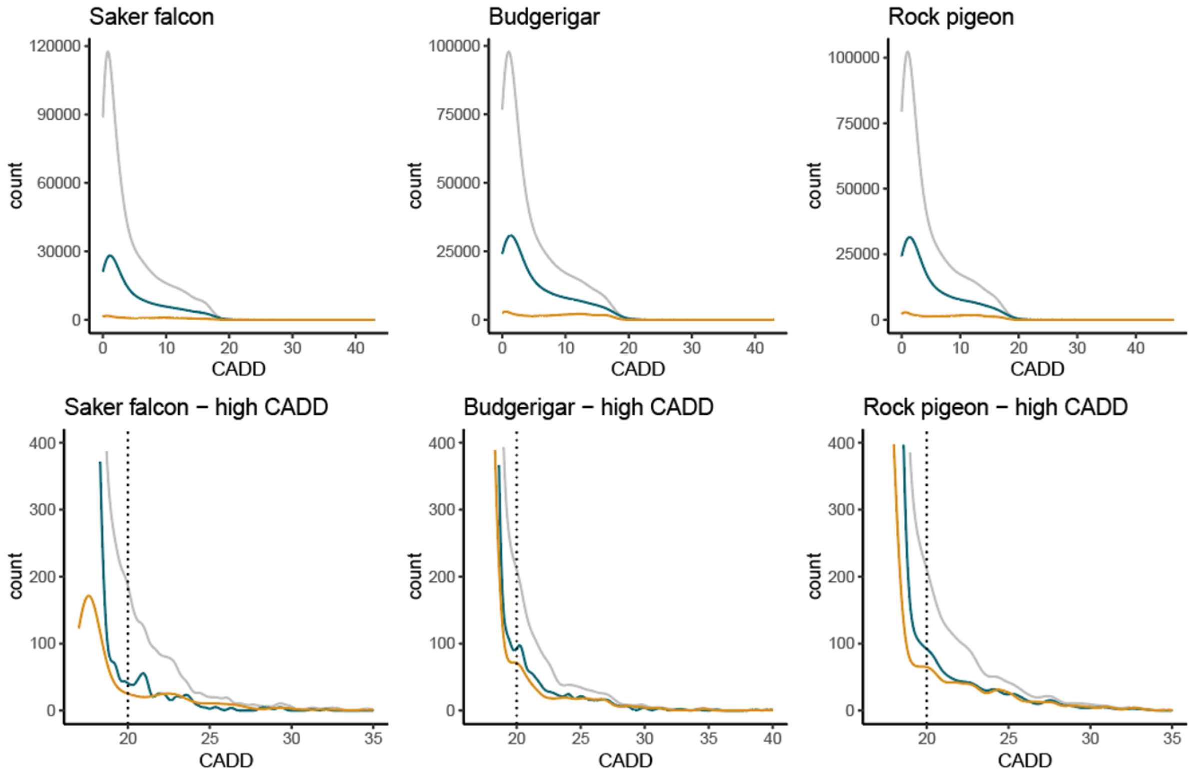


**Figure S3**. The count distributions of CADD score of homozygous substitutions to chicken (grey), filtering out sites shared by 20 or more species(data used in the main analyses, green) and the tests which compare to the ancestral node (method B, orange). After filtering out the sites where the ancestral node is different from chicken, only 4% of the substitutions previously identified compared to the chicken remained. On the other hand, 40% of the sites with CADD>20 passed this filter, a proportion roughly similar to the one obtained from filtering out shared sites by 20 or more species.

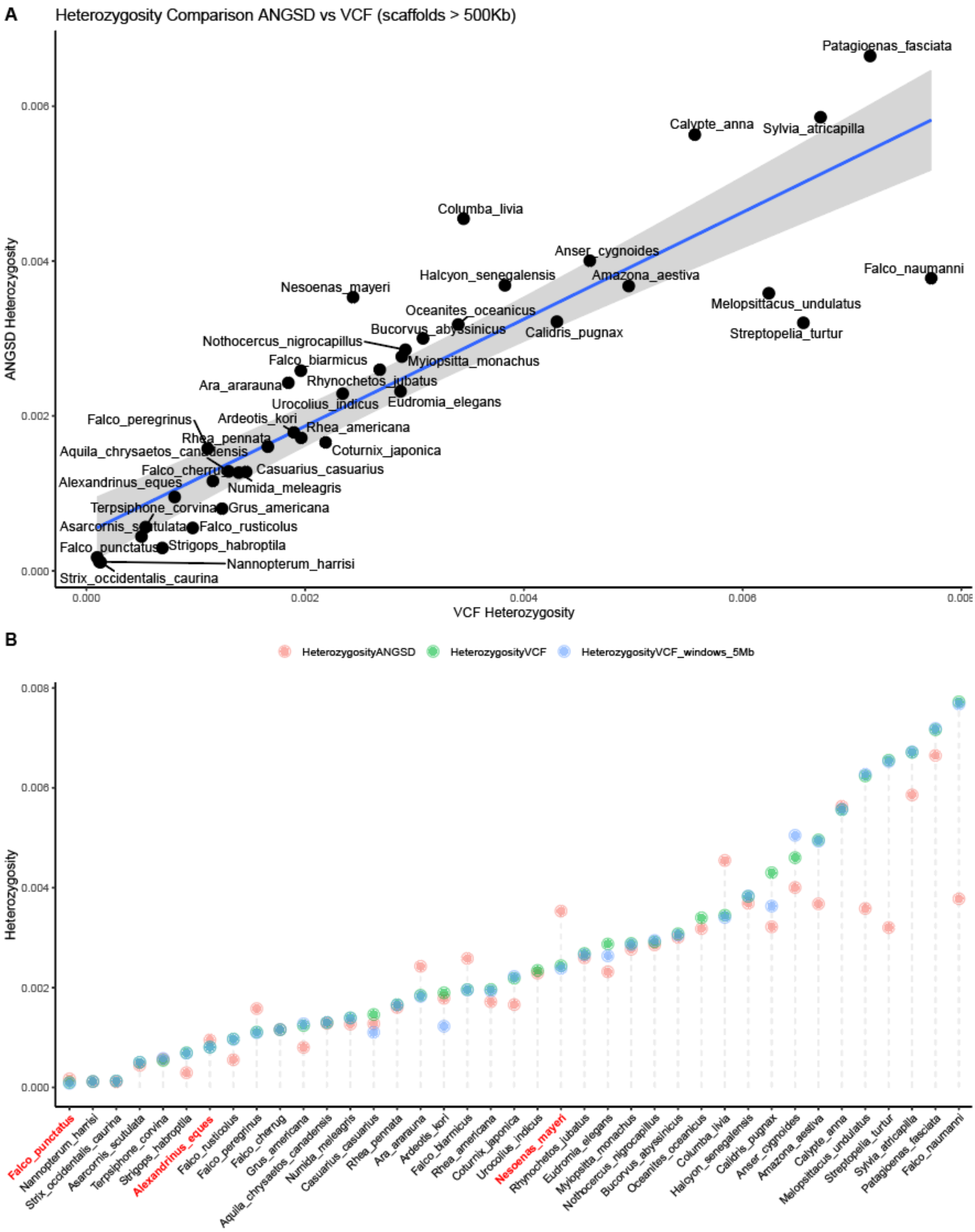


**Figure S4**. Heterozygosity comparison on scaffolds larger than 500Kb between ANGSD and counts from the VCF file. **A** Correlation between both estimates (Adjusted R-squared: 0.7798; p-value: 6.178e-14). **B** Heterozygosity values per sample calculated with ANGSD (HeterozygosityANGSD), and from the VCF in scaffolds > 500Kb (HeterozygosityVCF) and in scaffolds >5Mb (HeterozygosityVCF_windows_5Mb).

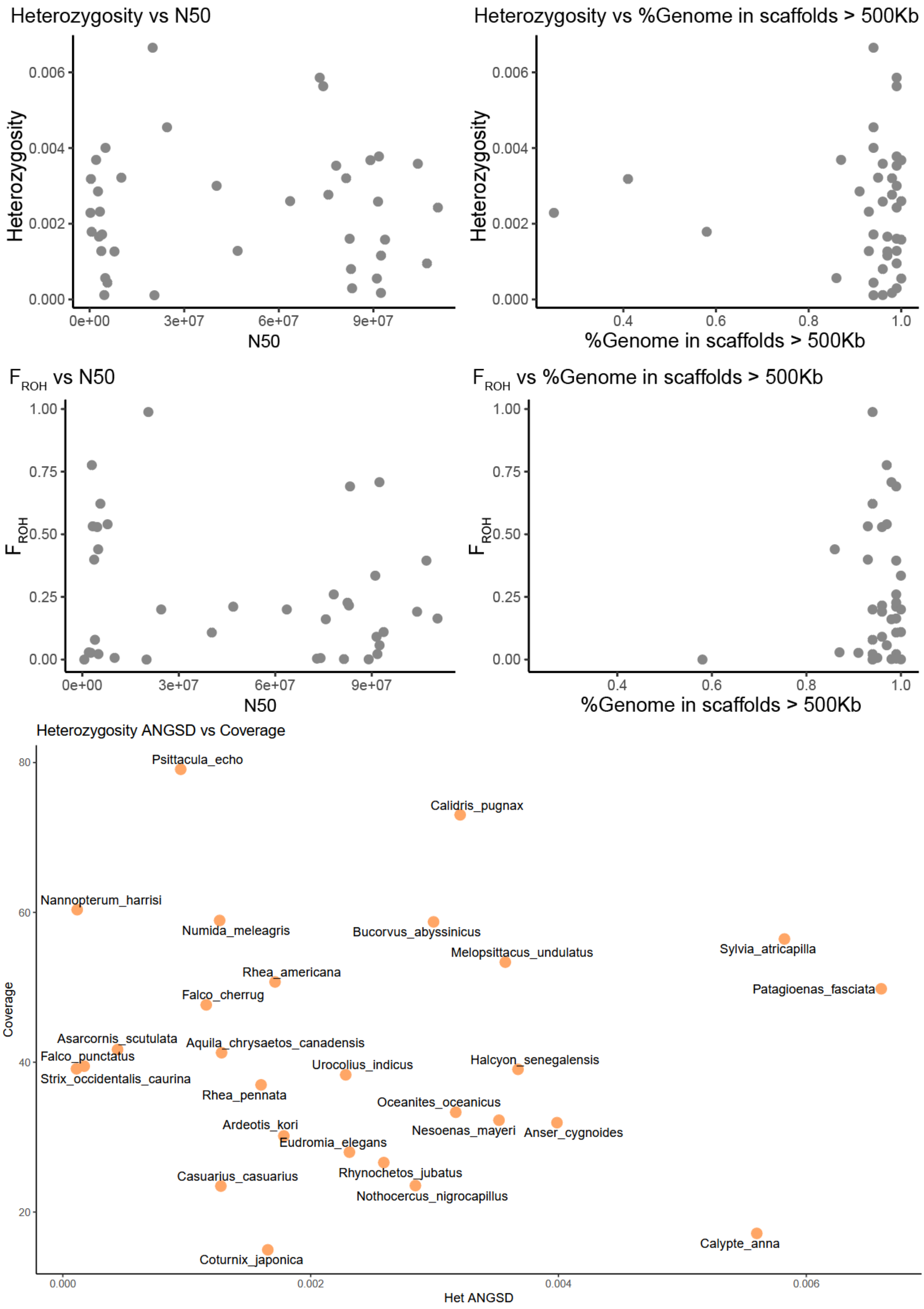


**Figure S5**. The quality of the reference genomes, measured in N50 of the scaffolds and the proportion of the genome in scaffolds longer than 500Kb, or the depth of the genome, had no effect on the estimation of genome heterozygosity or $F_{ROH}$.

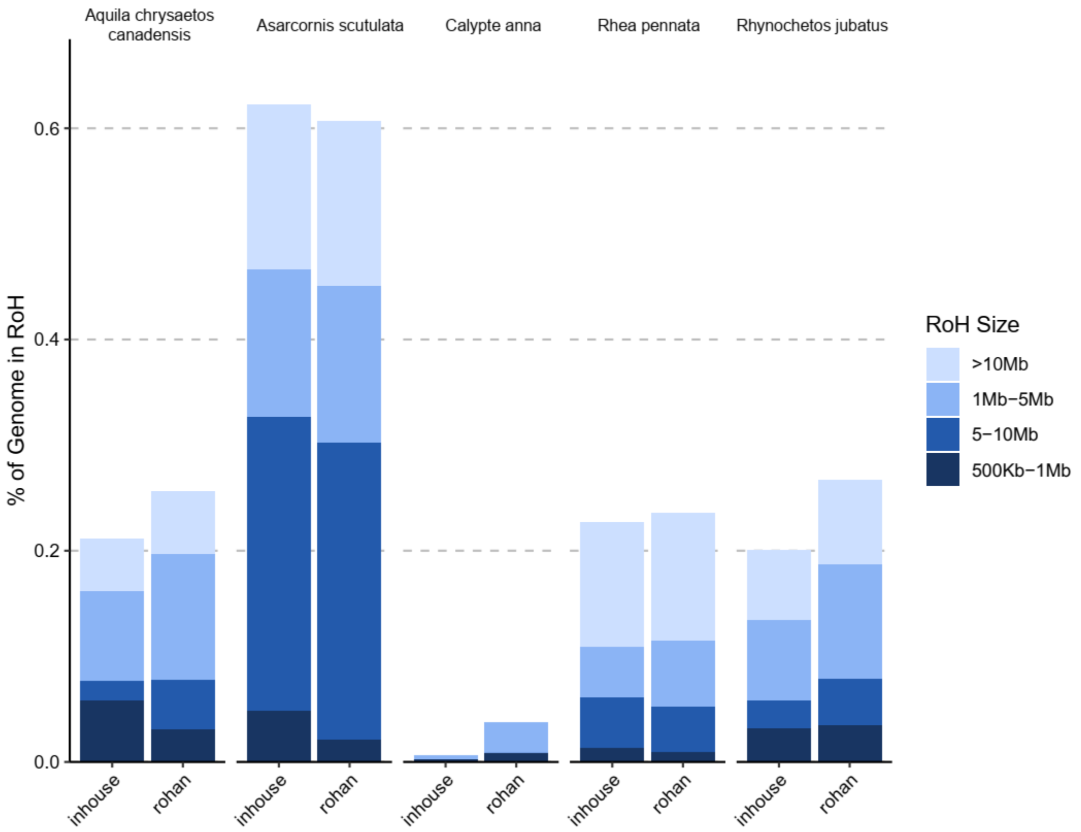


**Figure S6**. Comparison two estimators of runs of homozigosity, the inhouse method implemented in this work and rohan.

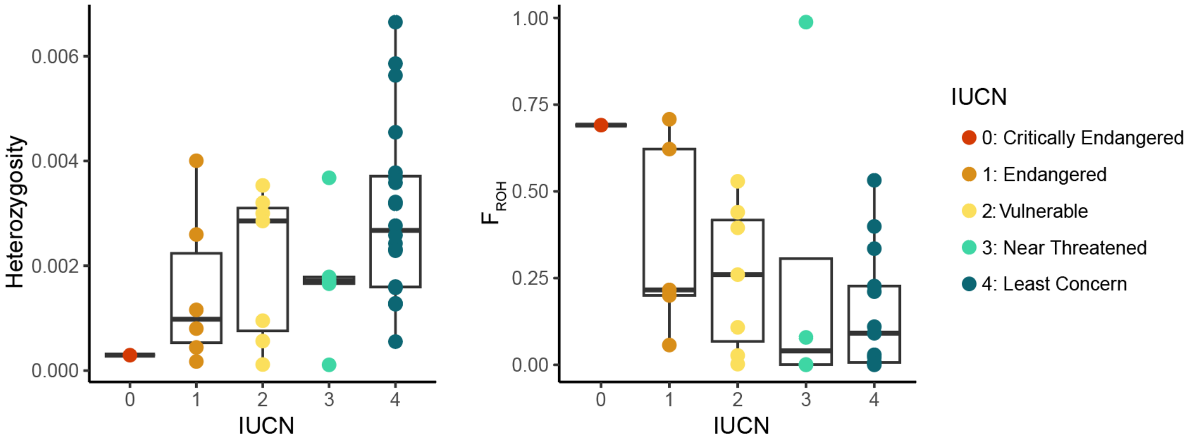


**Figure S7**. Heterozygosity and $F_{ROH}$ are not significantly correlated with IUCN Red List categories. GLM test for heterozygosity: $p$ = 0.07, $R^2$ = 0.13; for $F_{ROH}$: $p$ = 0.23, $R^2$ = 0.06.

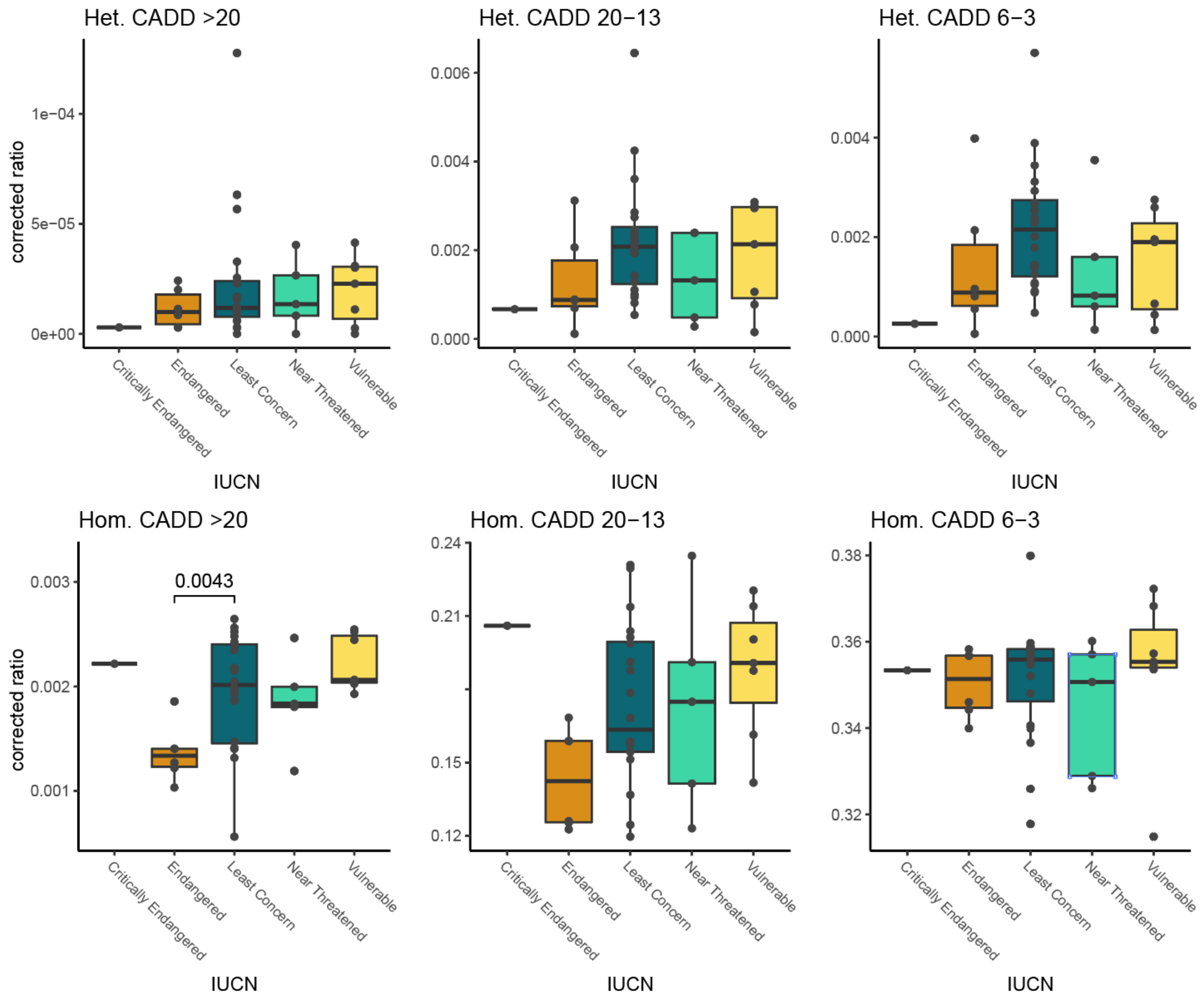


**Figure S8**. Heterozygous load (Het.) and putatively homozygous load (Hom.) in different IUCN Red List categories. Wilcoxon two-sample tests were performed between Least Concern and the rest categories, and only p values <0.05 are shown.

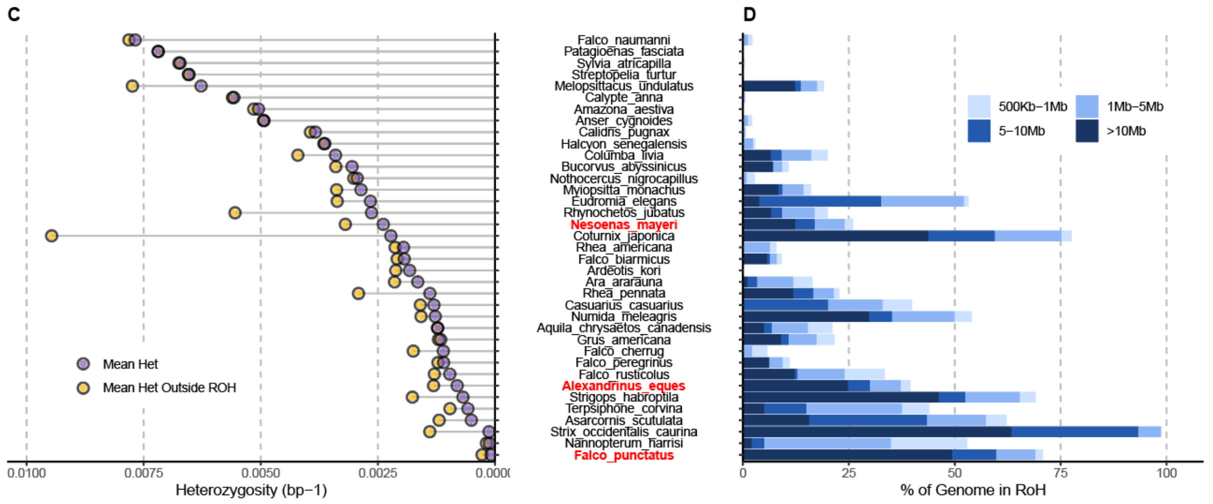


**Figure S9**. Relationship between heterozygosity and Runs of Homozygosity (ROH). **A)** Genome-wide heterozygosity (scaffolds >5Mb) and heterozygosity outside ROH. **B)** Percentage of the genome in ROHs (Inbreeding coefficient $F_{ROH}$).

**Table S1**. Metadata of the reference genomes and the species. Generation times and consensus population sizes (Nc) were collected from IUCN (https://www.iucnredlist.org/). Captive information was collected from the BioSample information of each reference genome. Mutation rates were from the species or closely related species and converted from mutation rates per year to mutation rates per generation using the generation times when data is unavailable.

**Table S2**. Genetic data estimated from alignments conducted in the study. “timetree_distance_chicken” refers to the divergence time of each species to chicken collected from Timetree of Life (http://timetree.org/); “hom_3-” refers to the raw counts of homozygous substitutions in UCEs with CADD score <3; “p_het_20+” refers to corrected ratio of heterozygous sites in UCEs with CADD score >=20; “p_hom_10-6” refers to corrected ratio of homozygous substitutions in UCEs with CADD score >=6 and <10, etc.

**Table S3**. PGLMM results for homozygous load models
